## Supplementary Figures for "Machine-learning predicts genomic determinants of meiosis-driven structural variation in a eukaryotic pathogen"

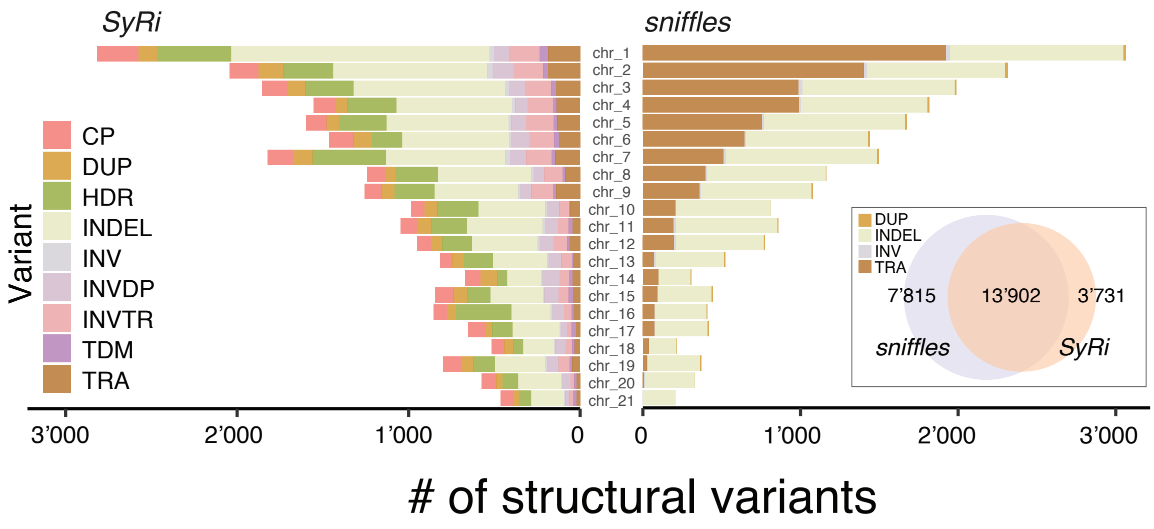


**Figure S1:** Number of structural variants identified on each chromosome using either the whole-genome alignment (SyRi) or the read-mapping method (sniffles) to the IPO323 reference genome. Both methods included a consolidation step for similar variants separated by less than 1,000 bp. Only translocations (TRA), indels (INDEL), inversions (INV) and duplications (DUP) are resolved using the sniffles method, while SyRi enabled the identification of copy variation (CP), highly diverged regions (HDR), inverted duplications (INVDP), inverted translocations (INVTR) and tandem repeats (TDM). The Venn diagram depicts the number of translocations, indels, duplications and inversions that were identified by both methods with overlapping positions.

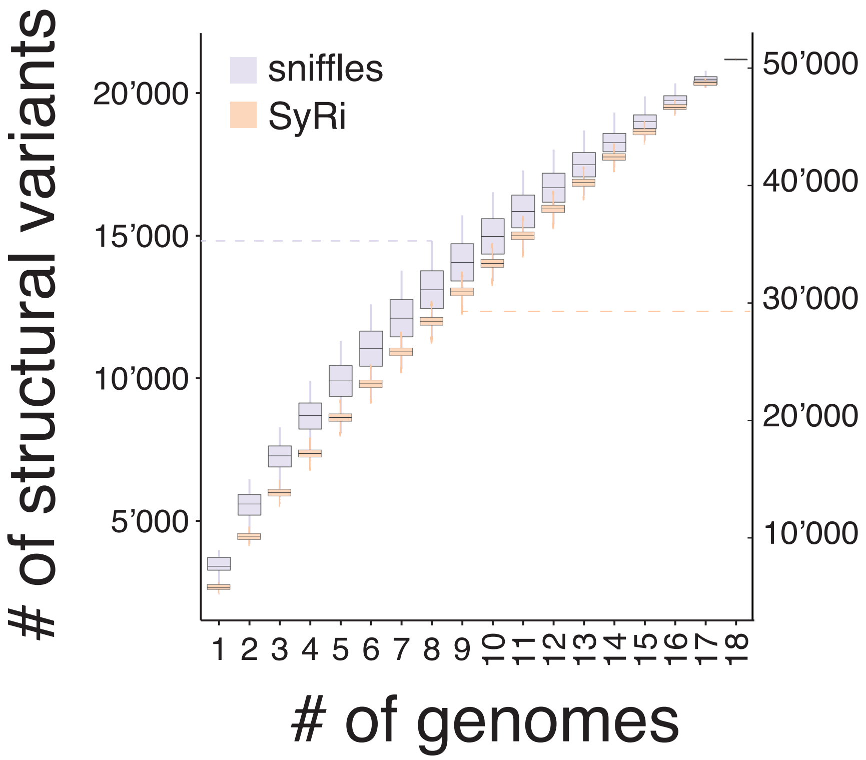


**Figure S2:** Number of structural variants recovered after resampling over 18 genomes. Results are shown for variants identified using either the whole-genome alignment (SyRi; y-axis right side) or the read-mapping method (sniffles; y-axis left side) to the IPO323 reference genome.

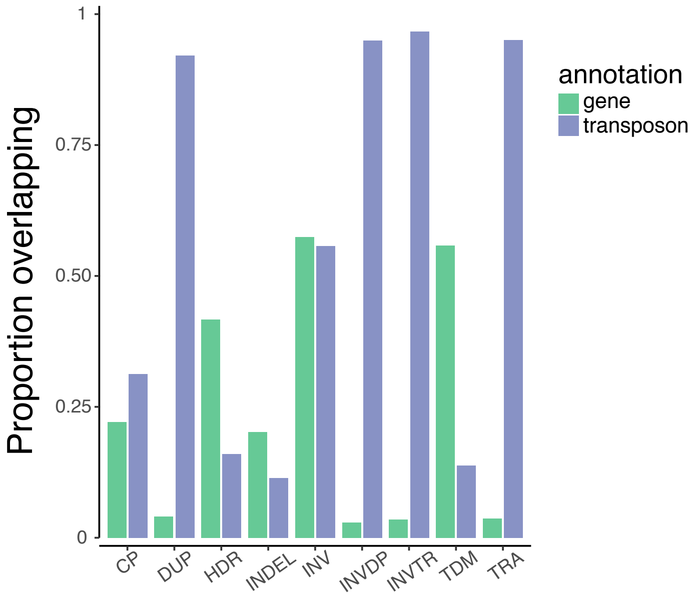


**Figure S3:** Proportion of translocations (TRA), indels (INDEL), inversions (INV), inverted duplications (INVDP), inverted translocations (INVTR), copy variation (CP), highly diverged regions (HDR), tandem repeats and duplications (DUP) overlapping with genes and transposable elements.

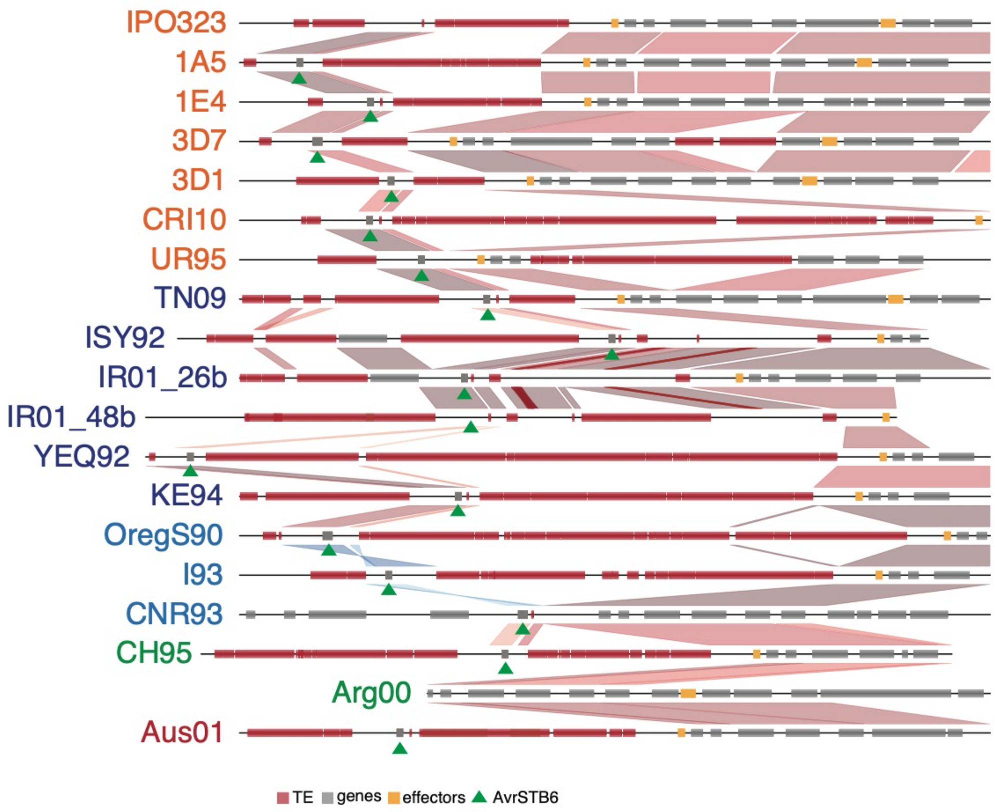


**Figure S4:** Synteny plot of the subtelomeric arm of chromosome 5 where the effector gene *AvrStb6* is located (denoted by the green triangles). The 19 *Zymoseptoria tritici* isolates are coloured as per their location of origin as in Figure 1A.

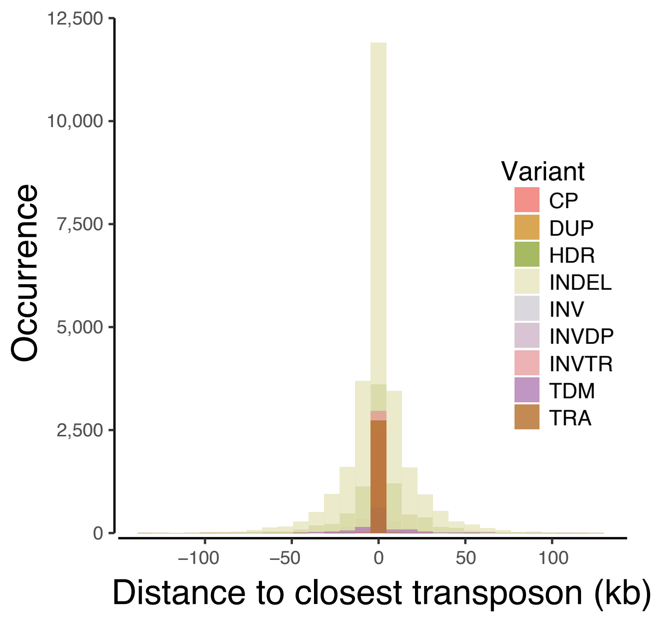


**Figure S5:** Occurrence of translocations (TRA), indels (INDEL), inversions (INV), inverted duplications (INVDP), inverted translocations (INVTR), copy variation (CP), highly diverged regions (HDR), tandem repeats and duplications (DUP) according to the distance to the closest transposable element. The distance is shown in kb.

| SV | model | Accuracy | Precision | Recall | F1 |
|---|---|---|---|---|---|
| CP | RF | 0.63 | 0.53 | 0.48 | 0.50 |
| DUP | RF | 0.88 | 0.67 | 0.84 | 0.74 |
| HDR | RF | 0.65 | 0.70 | 0.75 | 0.72 |
| INDEL | RF | 0.93 | 0.93 | 1.00 | 0.96 |
| INV | RF | 0.89 | NA | 0 | NA |
| INVTR | RF | 0.91 | 0.81 | 0.90 | 0.85 |
| INVDP | RF | 0.88 | 0.68 | 0.85 | 0.75 |
| TRA | RF | 0.91 | 0.82 | 0.88 | 0.85 |
| TDM | RF | 0.92 | 0 | 0 | NaN |
| CP | GLM | 0.61 | 0.51 | 0.51 | 0.51 |
| DUP | GLM | 0.88 | 0.68 | 0.80 | 0.74 |
| HDR | GLM | 0.65 | 0.71 | 0.74 | 0.72 |
| INDEL | GLM | 0.93 | 0.93 | 1.00 | 0.96 |
| INV | GLM | 0.89 | 0.22 | 0.02 | 0.04 |
| INVTR | GLM | 0.92 | 0.85 | 0.86 | 0.85 |
| INVDP | GLM | 0.90 | 0.72 | 0.82 | 0.77 |
| TRA | GLM | 0.91 | 0.88 | 0.82 | 0.84 |
| TDM | GLM | 0.92 | 0.14 | 0.02 | 0.03 |
| CP | ADA | 0.61 | 0.51 | 0.47 | 0.49 |
| DUP | ADA | 0.88 | 0.67 | 0.83 | 0.74 |
| HDR | ADA | 0.68 | 0.73 | 0.76 | 0.75 |
| INDEL | ADA | 0.93 | 0.93 | 1.00 | 0.96 |
| INV | ADA | 0.89 | NA | 0 | NA |
| INVTR | ADA | 0.91 | 0.81 | 0.90 | 0.85 |
| INVDP | ADA | 0.88 | 0.67 | 0.85 | 0.75 |
| TRA | ADA | 0.91 | 0.81 | 0.90 | 0.85 |
| TDM | ADA | 0.92 | 0 | 0 | NaN |
| CP | GBM | 0.61 | 0.50 | 0.52 | 0.51 |
| DUP | GBM | 0.87 | 0.65 | 0.85 | 0.74 |
| HDR | GBM | 0.68 | 0.72 | 0.78 | 0.75 |
| INDEL | GBM | 0.93 | 0.93 | 1.00 | 0.96 |
| INV | GBM | 0.89 | NA | 0 | NA |
| INVTR | GBM | 0.91 | 0.81 | 0.90 | 0.85 |
| INVDP | GBM | 0.88 | 0.67 | 0.86 | 0.75 |
| TRA | GBM | 0.91 | 0.81 | 0.89 | 0.85 |
| TDM | GBM | 0.92 | 0 | 0 | NaN |

**Figure S6:** Summary statistics of the models trained to predict translocations (TRA), indels (INDEL), inversions (INV), inverted duplications (INVDP), inverted translocations (INVTR), copy variation (CP), highly diverged regions (HDR), tandem repeats and duplications (DUP) when applied to the pangenome test dataset. For each type of structural variant, four models were trained using random forest (RF), logistic regression (GLM), boosted classification tree (ADA) and stochastic gradient boosting (GBM) algorithms.

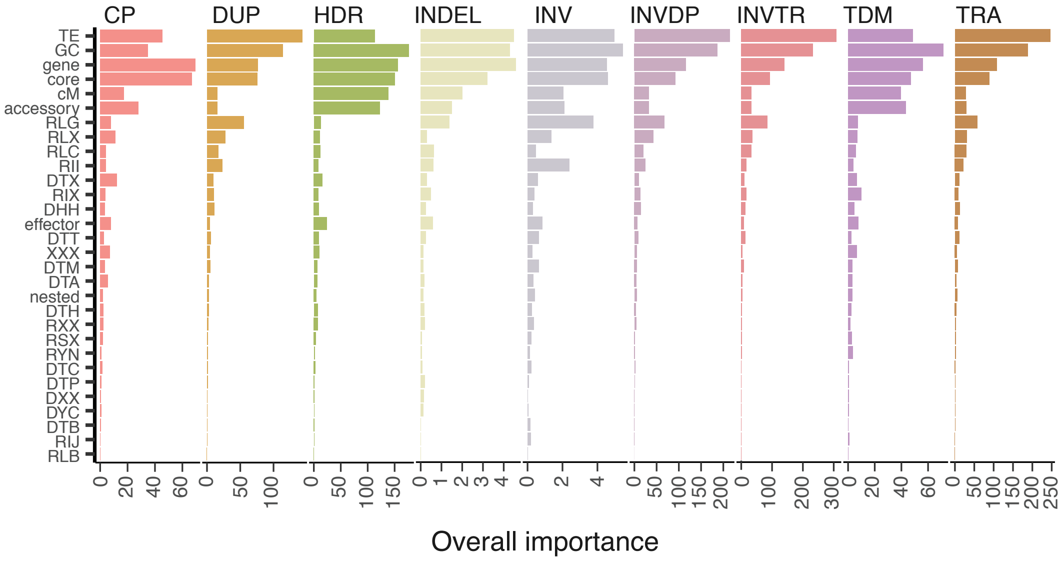


**Figure S7:** Relative importance of the sequence-based metrics used for modelling the occurrence of the nine types of structural variants using the random forest algorithm. Translocations (TRA), indels (INDEL), inversions (INV), inverted duplications (INVDP), inverted translocations (INVTR), copy variation (CP), highly diverged regions (HDR), tandem repeats and duplications (DUP)

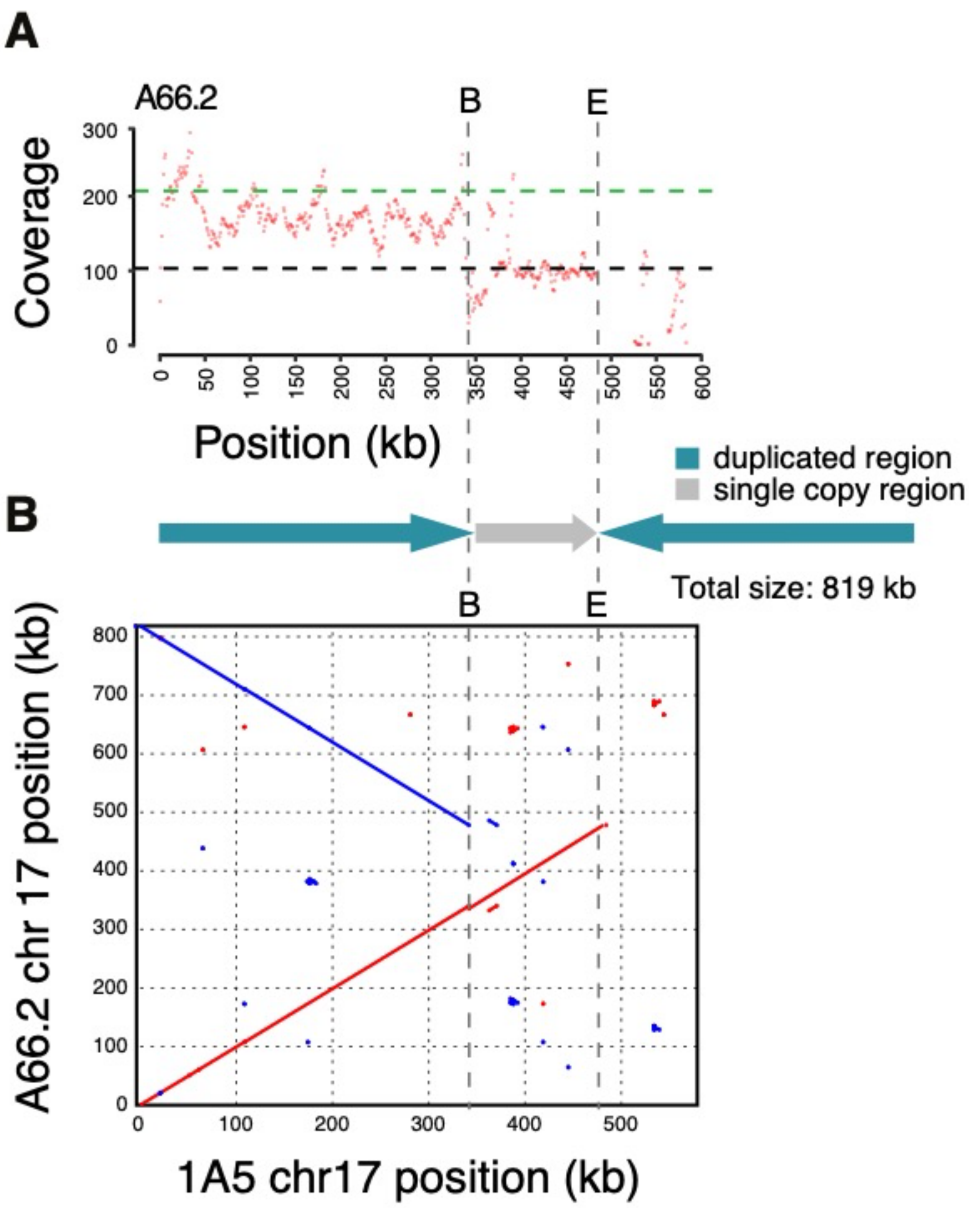


**Figure S8: A.** The coverage and breakpoints of the A66.2 progeny long-reads mapped to the 1A5 parent. Horizontal dashed lines indicate the mean of core chromosome coverage in black and in green the 2-fold mean of core chromosomes coverage. Red dots indicate the mean coverage in 1 kb windows (regions with >300X coverage were removed). Vertical dashed lines at B and E indicate positions where split reads map. **B.** Dotplots of the assembled chromosome 17 compared to the 1A5 parental chromosome. Inverted regions are indicated in blue.

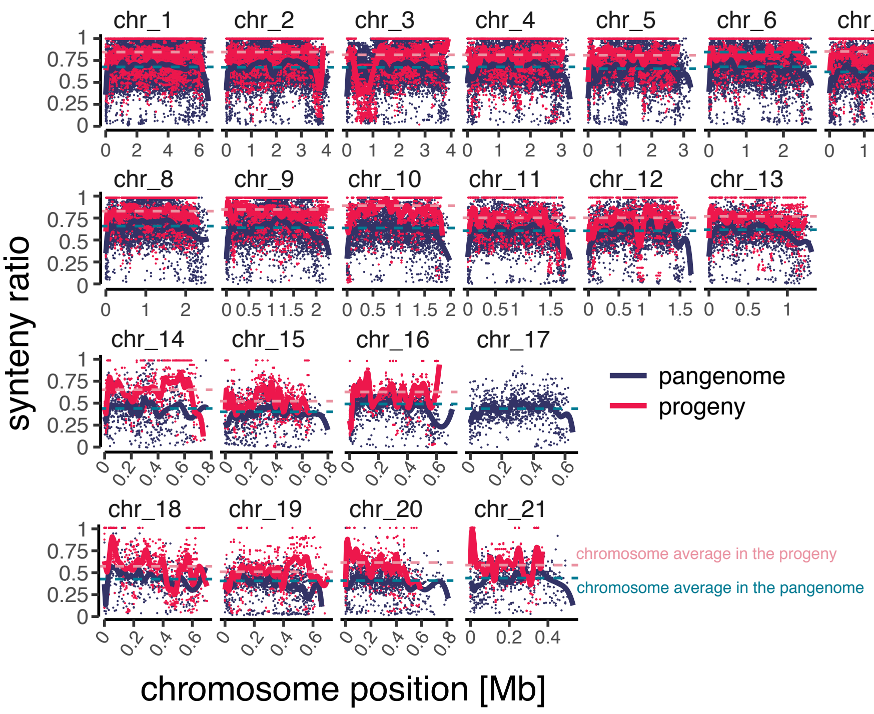


**Figure S9:** Genome-wide synteny expressed as a ratio (see Methods) across non-overlapping 10 kb windows in the progeny (in red) and compared to the pangenome (in blue). Coloured curves represent polynomial splines fitting calculated in R using the B-Splines basis with 25 degrees of freedom. Synteny of chromosome 17 is not shown for the progeny as it underwent multiple aberrant non-disjunction events making analyses of collinearity challenging.

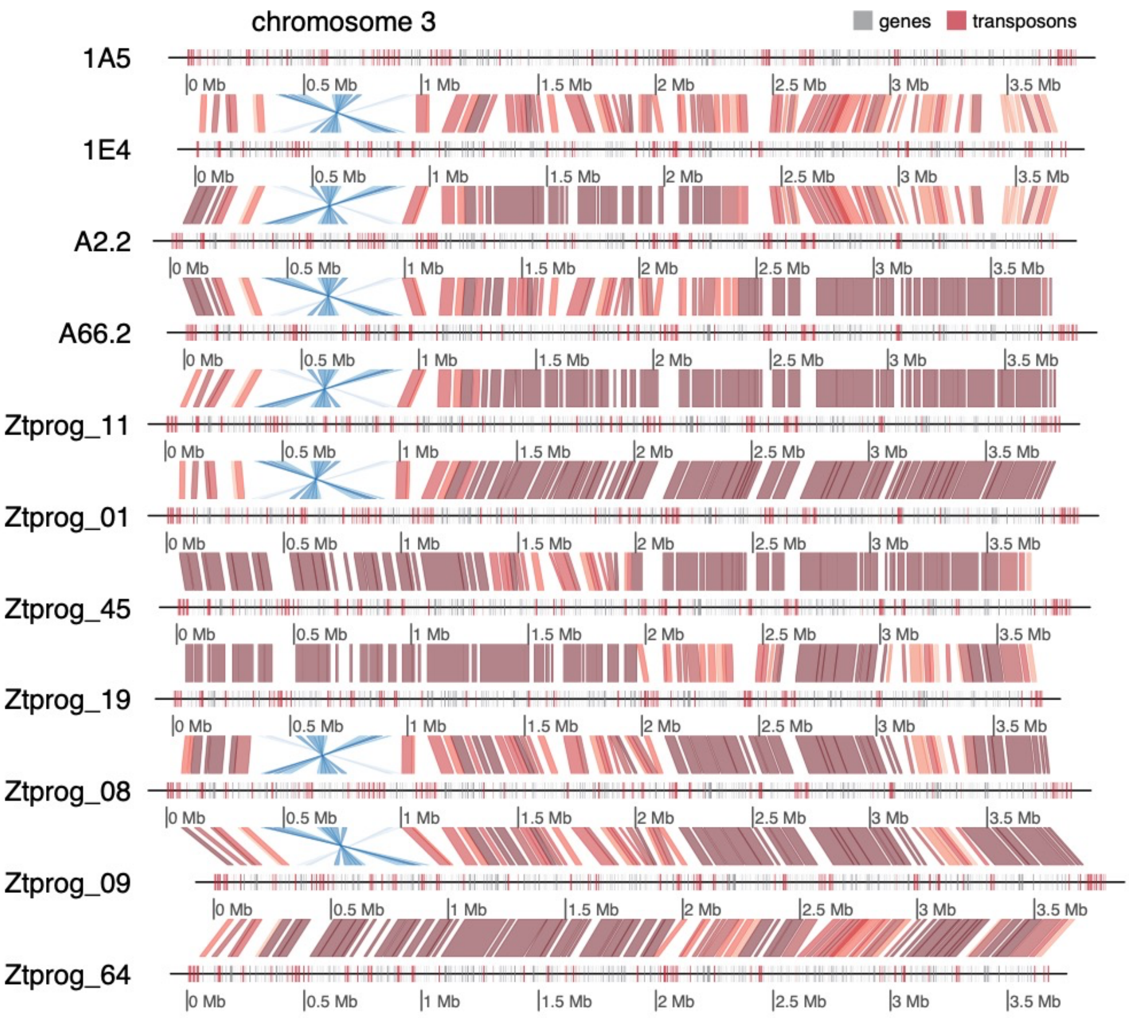


**Figure S10:** Synteny plot of chromosome 3 of different progeny in the pedigree depicting a large segmental inversion in a sub-telomeric region. The region shares low synteny in both the pangenome and the progeny pedigree (see Figure S3).

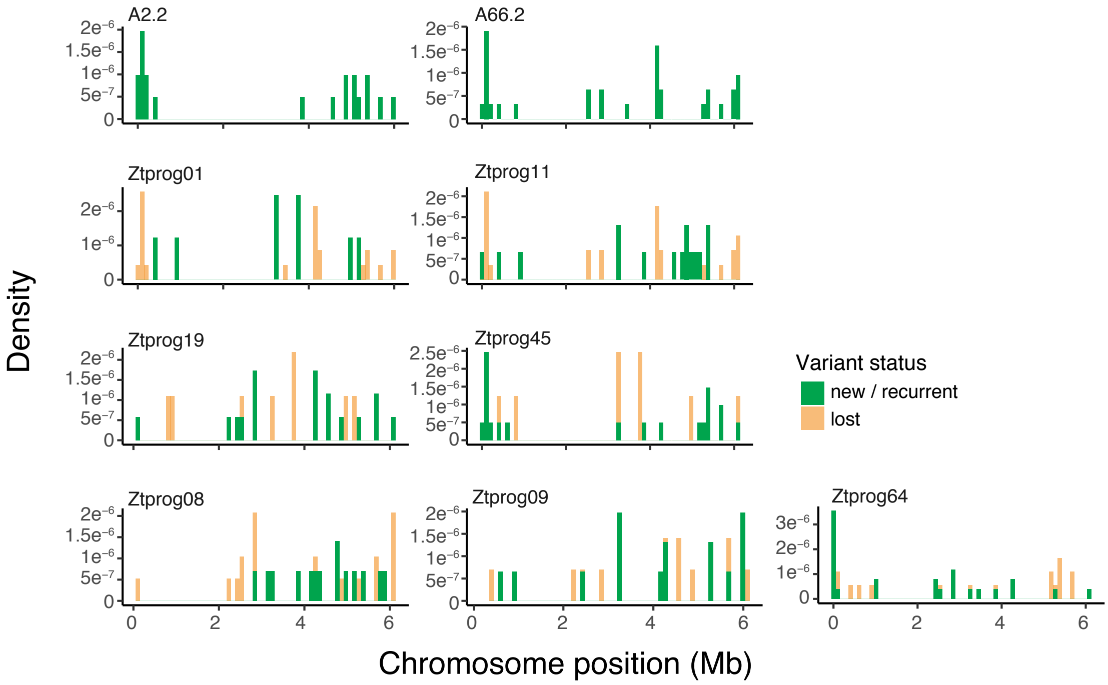


**Figure S11:** Density plot showing the distribution of new and recurrent structural variants compared to lost structural variants on chromosome 1 as observed in the pedigree.

| SV | model | Accuracy | Precision | Recall | F1 |
|---|---|---|---|---|---|
| CP | RF | 0.64 | 0.04 | 0.78 | 0.08 |
| DUP | RF | 0.80 | 0.18 | 0.79 | 0.29 |
| HDR | RF | 0.38 | 0.03 | 0.74 | 0.05 |
| INDEL | RF | 0.05 | 0.05 | 1.00 | 0.09 |
| INV | RF | 1.00 | NA | 0.00 | NA |
| INVTR | RF | 0.73 | 0.08 | 0.89 | 0.14 |
| INVDP | RF | 0.82 | 0.23 | 0.81 | 0.36 |
| TRA | RF | 0.73 | 0.06 | 0.94 | 0.11 |
| TDM | RF | 0.98 | 0.09 | 0.09 | 0.09 |
| CP | GLM | 0.64 | 0.04 | 0.75 | 0.07 |
| DUP | GLM | 0.82 | 0.19 | 0.75 | 0.30 |
| HDR | GLM | 0.38 | 0.03 | 0.74 | 0.05 |
| INDEL | GLM | 0.05 | 0.05 | 0.99 | 0.09 |
| INV | GLM | 0.99 | 0.00 | 0.00 | NaN |
| INVTR | GLM | 0.76 | 0.08 | 0.82 | 0.15 |
| INVDP | GLM | 0.84 | 0.24 | 0.74 | 0.37 |
| TRA | GLM | 0.78 | 0.06 | 0.87 | 0.12 |
| TDM | GLM | 0.99 | 0.07 | 0.06 | 0.07 |
| CP | ADA | 0.64 | 0.04 | 0.79 | 0.08 |
| DUP | ADA | 0.80 | 0.18 | 0.79 | 0.29 |
| HDR | ADA | 0.38 | 0.03 | 0.79 | 0.05 |
| INDEL | ADA | 0.05 | 0.05 | 1.00 | 0.09 |
| INV | ADA | 1.00 | NA | 0.00 | NA |
| INVTR | ADA | 0.73 | 0.07 | 0.88 | 0.14 |
| INVDP | ADA | 0.81 | 0.23 | 0.81 | 0.35 |
| TRA | ADA | 0.71 | 0.05 | 0.96 | 0.10 |
| TDM | ADA | 0.98 | 0.06 | 0.06 | 0.06 |
| CP | GBM | 0.61 | 0.04 | 0.79 | 0.07 |
| DUP | GBM | 0.79 | 0.18 | 0.83 | 0.29 |
| HDR | GBM | 0.35 | 0.03 | 0.80 | 0.05 |
| INDEL | GBM | 0.05 | 0.05 | 1.00 | 0.09 |
| INV | GBM | 1.00 | NA | 0.00 | NA |
| INVTR | GBM | 0.72 | 0.07 | 0.88 | 0.14 |
| INVDP | GBM | 0.81 | 0.23 | 0.82 | 0.35 |
| TRA | GBM | 0.72 | 0.06 | 0.96 | 0.11 |
| TDM | GBM | 0.98 | 0.09 | 0.15 | 0.11 |

**Figure S12:** Summary statistics of the models trained to predict translocations (TRA), indels (INDEL), inversions (INV), inverted duplications (INVDP), inverted translocations (INVTR), copy variation (CP), highly diverged regions (HDR), tandem repeats and duplications (DUP) when applied to the progeny dataset. For each type of structural variant, we show the results of the four models trained using random forest (RF), logistic regression (GLM), boosted classification tree (ADA) and stochastic gradient boosting (GBM) algorithms.

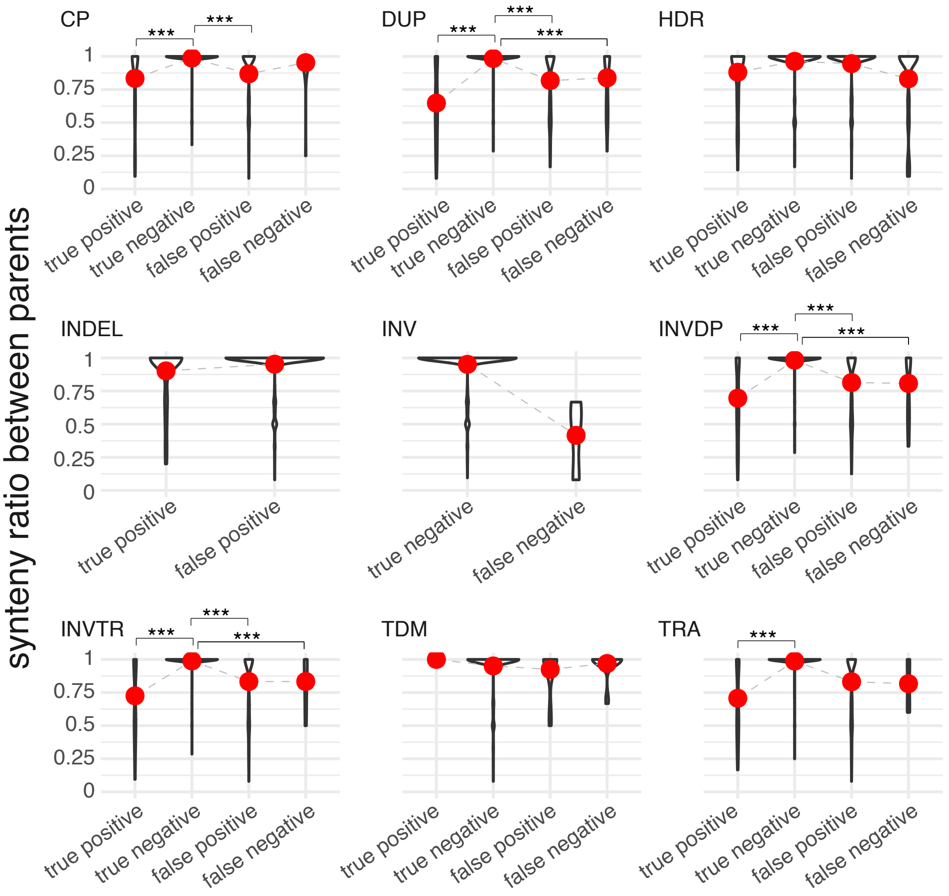


**Figure S13:** Violin plots depicting the distribution of synteny values in 10 kb windows for accurate model predictions (true positives and true negatives) and wrong model prediction (false negatives and false positives). Values for each type of rearrangement predictions are shown separately. Asterisks show significant differences ($p$ < 1e$^{-6}$) based on the Wilcoxon rank-sum test.

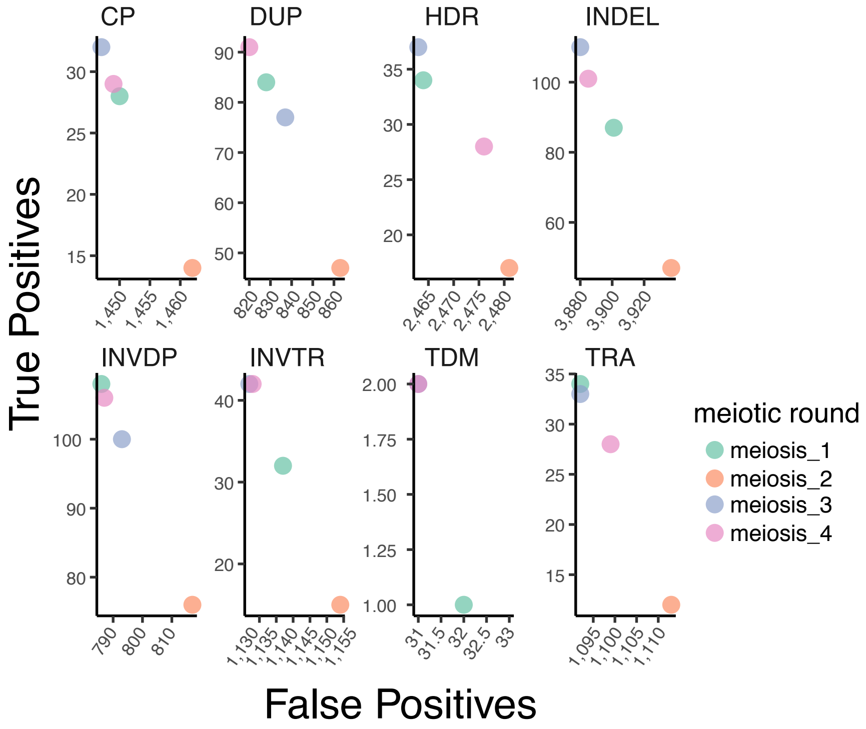


**Figure S14:** Model performance for each progeny set of structural variants generated in the parental round of meiosis (noted M1 to M4, for meiotic round 1 to 4). Performance is shown as the relationship between the true positive rate over the false positive rate. M1 regroups structural variants identified in the progeny A2.2 and A66.2, M2 regroups structural variations identified in progeny Ztprog_01 and Ztprog_11, M3 regroups structural variations identified in progeny Ztprog_19 and Ztprog_45 and M4 regroups structural variations identified in progeny Ztprog_08, Ztprog_09 and Ztprog_64. Results for each type of variant are shown separately.

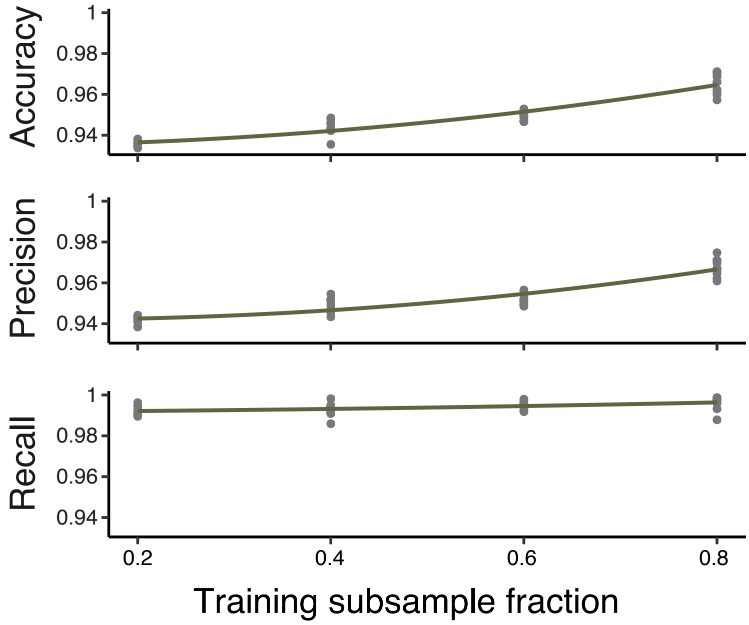


**Figure S15:** Variation of performance metrics for the random forest model trained on 10 random subsamples representing a fraction of 0.2, 0.4, 0.6 and 0.8 of the total pangenome dataset.
